# Fitness Estimation for Genetic Evolution of Bacterial Populations

**DOI:** 10.1101/2020.10.03.324830

**Authors:** Sergey S. Sarkisov, Ilya Timofeyev, Robert Azencott

## Abstract

In this paper we develop and test algorithmic techniques to estimate genotypes fitnesses by analysis of observed daily frequency data monitoring the long-term evolution of bacterial populations. In particular, we develop a non-linear least squares approach to estimate selective advantages of emerging new mutant strains in locked-box stochastic models describing bacterial genetic evolution similar to the celebrated Lenski experiment on Escherichia Coli. Our algorithm first analyses emergence of new mutant strains for each individual trajectory. For each trajectory our analysis is progressive in time, and successively focuses on the first mutation event before analyzing the second mutation event. The basic principle applied here is to minimize (for each trajectory) the mean squared errors of prediction *w*(*t*) − *W*(*t*) where the observed white cell frequencies *w*(*t*) are predicted by *W*(*t*), which is computed as the conditional expectation of *w*(*t*) given the available information at time (*t* − 1). The pooling of all selective advantages estimates across all trajectories provides histograms on which we perform a precise peak analysis to compute final estimates of selective advantages. We validate our approach using ensembles of simulated trajectories.

## 1 Introduction

Studies of evolutionary dynamics, including bacterial populations, received a considerable amount of attention in recent decades starting with the celebrated Lenski experiment [15, 26]. Since then other groups contributed to experimental studies of Escherichia Coli (see e.g. [3, 10, 2, 16, 6, 11, 4]). Typical setups for these experiments involve multiple populations processed in parallel. These populations undergo daily growth followed by daily selections of fixed-size samples. Initially, all populations consist only of cells having the same “ancestor” genotype. Since most evolutionary experiments are carried out for a long time, a varied range of random mutations occur in different populations, so that the genetic composition of different populations can differ drastically over time. The main goal of such experiments is to understand major features of evolutionary dynamics, including the rate of evolutionary change, likelihood of emergence of a new genoype, fitness landscape, probabilities of fixation for emerging mutants, etc.

The Lenski long-term evolution experiments on Escherichia coli provided a considerable insight and evidence for the mechanisms of growth and adaptation of asexual organisms (e.g. [15, 26, 8]). Analysis of the experimental data shed some light upon the relative growth rates of stronger mutants and their ancestors. The Lenski experimenst primarily focuses on the overall population adaptation [1, 27]. Our goal is to develop and test an efficient algorithm dedicated to estimating individual selective advantages for emerging mutants.

The adaptive evolution of bacterial populations is driven by a random emergence of beneficial mutations and spread of the fittest mutants due to natural random selection. Various mathematical models of this evolution process have been developed (see e.g. [12, 25]), and estimation of process parameters in these models is always one of the key issues. The evolutionary dynamics of bacterial populations depends on two family of parameters, namely the selective advantages of mutants and the rate of mutations [10, 17, 14]. In this paper we assume that the mutation rate *μ* is roughly the same for all emerging mutants, and is already known. Therefore, we focus on estimating selective advantages for multiple competing mutants. Our techniques can be easily extended to estimate μ as well. Several previous papers addressed similar questions. For instance, one study involved both experimental and theoretical analyses of evolving viral populations [21, 20]. These papers include estimation for their adaptation models parameters, as well as for the distribution of beneficial mutation effects. However, this work was limited to viruses, because of their small genome size, high mutation rate, and large mutation effects. By contrast, in bacterial populations, direct estimates of mutational parameters is much more complex [23] because these parameters are usually estimated indirectly by inference from observable markers [23, 10, 2]. A more recent study, involving Tim Cooper’s experiments on Escherichia Coli, focused on populations where a single mutant type emerges and overtakes the population [28]. Mathematical derivations and intensive model simulations were combined to derive accurate estimates for the mutation rate and selective advantage of the first mutant reaching fixation. Another earlier parameter estimation study [10] used non-linear regression to analyze the first emerging mutant. Our approach here is complementary to these earlier studies. In particular, we develop efficient parameter estimation algorithms to cover the emergence and competition of multiple mutant types.

In this paper, we consider mathematical setup motivated by the Escherichia Coli evolutionary experiments [28, 10, 2]. In such experiments one observes many populations simultaneously evolving in parallel. Each initial population consists entirely of *N* cells with the same “ancestor” genotype, with N typically ranging from 50,000 to 100,000. In each population, half of the cells are colored via a particular marker (e.g. Ara^+^) and the other half are colored via a different marker (e.g. Ara^−^). These two markers define “white” and “red” cell colonies, initially coexisting in proportions 50/50 in all populations. Cell dynamics is not altered by these markers. Asexual reproduction occurs via binary fission, and genetic markers are passed down from parents to offspring [2]. The populations are allowed to grow daily until exhaustion of the daily initial glucose or equivalent “food supply”, which happens in less than 24 hours. At that point each cell population reaches a size in the range of 10^7^ to 10^8^, and a sub-population of approximate size N is extracted by dilution and transferred to a new culture of fresh growth medium. This transfer step is repeated daily for all populations. The white and red cell frequencies are estimated periodically for instance by plating cells on indicator media. Therefore, after initialization of all populations with identical “ancestor” genotype and 50/50 marker coloration into “white” and “red” colonies, the daily cycle of growth, mutations, and dilution is repeated for hundreds of days. A beneficial mutation emerging on day *t* among white cells can be detected at a later time *t + s* as soon as it generates a significant upward change in the frequency *w*(*t + s*) of white cells. Similarly beneficial mutations emerging among red cells at time *t* can be detected by significant downward change in *w*(*t + s*). Ideally the whole trajectory *w*(*t*) can be recorded for each population. After a rough detection for the emergence of new mutant strains, we apply a non-linear least squares method to estimate more accurately the times of emergence of these new strains, as well as their selective advantages. This is done separately for each trajectory. We then combine estimated selective advantages computed for each individual trajectory. By a detailed analysis of the histogram of all these estimates, we finalize more accurate estimates for the selective advantages of the observed mutant strains. Our stochastic model for the evolutionary dynamics of bacterial populations is a Markov chain where the cycle of growth, mutation, selection is repeated daily. To implement our non-linear least squares estimation of model’s parameters, we develop explicitly computable predictors of next day white cell frequency, and minimize the mean squared errors of these predictors with respect to the observed data. We validate our approach on simulated data. In particular, we use Markov chain model to numerically generate an ensemble of evolutionary trajectories. We then apply our estimation algorithm to these simulated trajectories and evaluate accuracy of our estimated selective advantages. We utilize Gaussian mixture approach to analyze numerical results, which validate the statistical consistency of our parameters estimators.

## 2 Stochastic Model

Various modifications of Markov chain “locked-box” models have been often used to describe evolutionary experiments outlined above (e.g. [28, 19, 10, 22, 25]). In these models a daily random mutation phase occurs right after a daily deterministic growth phase, and is followed by a random selection phase, formalized by extraction of a random sample of fixed size *N*. The evolutionary experiment is then modeled by repeating the daily cycle *growth* ⟹ *mutations* ⟹ *selection* ⟹ ….

We consider only a finite number *i* ∈ {1,2,3,…, *g*} of possible genotypes. The ancestor genotype is denoted as *i* = 1 and the other (*g* − 1) genotypes are mutants. Let *t* ∈ {1,2,…} be the discrete time index (measured in days). Denote *h_i_*(*t*), the frequency of genotype *i* at the beginning of day *t*. The random vector *h*(*t*) = {*h*_1_(*t*),…, *h_g_*(*t*)} clearly verifies 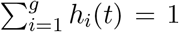, and is the whole histogram of genotypes frequencies at the beginning of day *t*. Next, we describe the three daily phases in more detail. First, we describe the Markov chain model for the evolution of a single-color population; extension to two colors is done in a rather straightforward manner by doubling the number of cell types.

### 2.1 Daily Growth Phase

At the beginning of any day *t*, the current cell population always has total size *N*, and is partitioned into *g* sub-populations *POP_i_*(*t*) consisting of cells with the same genotype *i*. Let *N_i_*(*t*) = *N* × *h_i_*(*t*) be the size of *POP_i_*(*t*), so that 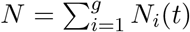. The daily growth phase is modeled by a deterministic exponential growth in which cell divisions occur at fixed exponential rate *ρ_i_* for all cells of genotype *i*. Therefore, at the end of the day *t* growth phase, which has fixed duration *D*, the size of *POP_i_*(*t*) becomes

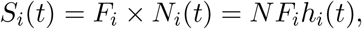

where the multiplicative daily growth factor *F_i_* = exp(*ρ_i_D*) is specific to genotype *i*. Hence, the total population size at the end of the day *t* growth phase is 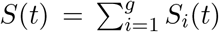. Without loss of generality, assume that genotypes are ordered with respect to their growth factors so that *F*_1_ < *F*_2_ < ⋯ < *F*_*g*−1_ < *F_g_*. The *selective advantage s_i_* of genotype *i* with respect to the ancestor genotype 1, is defined by

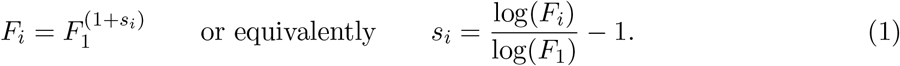

Some models (see e.g. [7, 28]) assume that the restricted daily amount of nutrients forces the daily population growth to stop when population reaches a fixed large maximum size. However, in this paper, our model assumes that the daily growth phase has a fixed duration *D*, already integrated in the multiplicative daily growth factors *F*_1_,…, *F_g_*. At the end of the day *t* growth phase one has *NF*_1_ ≤ *S*(*t*) ≤ *NF_g_*.

### 2.2 Daily Mutations Phase

At the end of the day *t* growth phase, the population size *S*(*t*) is typically several orders of magnitude larger than the daily initial population size, *N*. For instance, in Escherichia Coli long term experiments *N* ranges from 10^5^ to 10^8^, and *F*_1_ ≈ 200, while the mutation rate *μ* ranges from 10^−7^ to 10^−6^. Therefore, *S*(*t*) ranges from 10^7^ to 10^10^, and every day a positive number of new mutants can be expected to emerge during the daily mutation phase. Here we consider only beneficial mutations, since deleterious mutations are typically quickly eliminated from the population due to a lower selective advantage.

In our mathematical model, we assume that the daily mutations are independent random events, simultaneously occurring at the end of the daily growth phase. Next, we describe precise mathematical formulation for mutations.

At the end of the day *t* growth phase, and just before the day *t* mutations phase, the day *t* initial population *POP_i_*(*t*) of cells with genotype *i*, has grown into a much larger population *GPOP_i_*(*t*), of size *S_i_*(*t*). Denote *ν_i_ = ν_i_*(*t*) the random number of mutants emerging from population *GPOP_i_* during the mutations phase. We assume that the conditional distribution of *ν_i_*(*t*) given the initial day *t* histogram *h*(*t*) is a Poisson distribution with mean

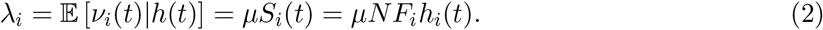

Note that the mutation rate *μ* is assumed to be the same for all genotypes. Our approach can be easily extended to mutation rates *μ_*i*_* which depend on genotype *i*, provided all *μ_i_* are small and are of the same order of magnitude. The conditional distribution of *ν_i_* is hence given by

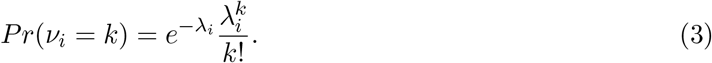

We assume that all *ν_i_*(*t*), with 1 ≤ *i* ≤ *g* are conditionally independent given the histogram *h*(*t*).

Our mutation model also involves a fixed (unknown) transition matrix 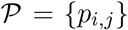 with 0 ≤ *p_i,j_* ≤ 1 and 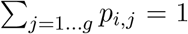. The matrix 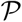 governs the stochastic transition from genotype *i* to genotype *j* by a single random mutation. Hence for each i, any mutant emerged from *GPOP_i_*(*t*) has probability *p_i,j_* of having genotype *j*. Of course one imposes *p_ii_* = 0 for all *i*.

Denote *ν_i,j_ = ν_i,j_*(*t*) the random number of mutants emerging from population *GPOP_i_*(*t*) and mutating into cells of genotype *j*. Then we have

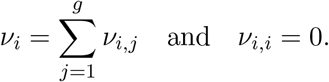

The conditional distribution of the random vector [*ν*_*i*,1_,…, *ν_i,g_*] given both *h*(*t*) and *ν_i_*(*t*) is then naturally assumed to be a *multinomial distribution* with probabilities {*p*_*i*,1_,…*p_i,g_*} within a population of size *ν_i_*(*t*). Hence for each *i* and for all non-negative integers *k*_1_,…, *k_g_* verifying *k_i_* = 0 and ∑_*j*=1…*g*_ *k_j_ = ν_i_*, we have

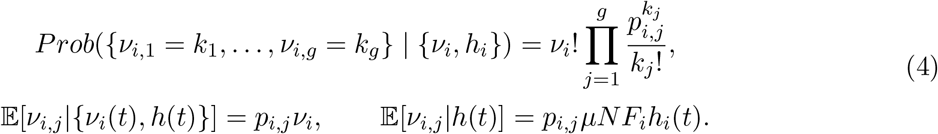

For any fixed genotype *i*, denote *EM_i_* and *IM_i_* the respective numbers of ‘emigrants” from population *GPOP_i_* and of ‘immigrants” into *GPOP_i_*. More precisely let

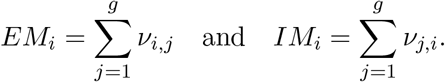

Let Δ*S_i_*(*t*) = *IM_i_ − EM_i_* be the net change in the size of population *GPOP_i_*. One can easily check that the mutation phase does not alter the total population size, i.e. 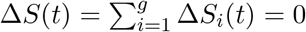.

### 2.3 Daily Selection Phase

On each day *t*, after the growth phase and the mutations phase, a *random selection* is performed by extracting a random sample of *fixed size N* from the whole terminal population on day *t*, which still has size *S*(*t*) >> *N*. In practical laboratory experiments, this selection is often implemented by culture dilution.

Denote *f_i_* the frequency of genotype *i* after growth and mutations. We clearly have

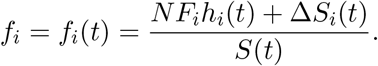

The daily random selection of *N* cells within a day *t* terminal population of size *S*(*t*) is then modelled by a multinomial distribution based on the probability vector *f*_1_,…, *f_g_*. In particular, let *κ_i_* be the number of genotype-*i* cells after the day *t* random selection. The joint distribution of *κ_i_, i* = 1,…, *g* is given by

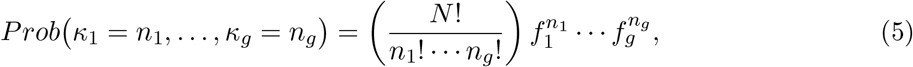

where 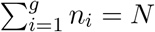.

The *N* cells randomly selected at the end of day *t* have a random histogram *h*(*t* + 1). After physical or virtual “transfer” these *N* cells initiate the day *t* + 1 cycle of growth / mutations / selection. Thus the day *t* + 1 cycle begins with a population of size *N* and histogram *h*(*t* + 1).

The cycle of {deterministic growth, random mutations, random selection} is repeated daily for the whole duration of the long-term experimental run. The process modeled by these successive three steps is a discrete-time Markov Chain driving the random vectors *h*(*t*) = (*h*_1_ (*t*),…, *h_g_*(*t*)) which describe the time-evolution of genotype frequencies. All frequencies sum up to one, i.e. 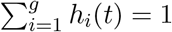, and a sample path of this process can be easily simulated given the vector of initial genotypes frequencies *h*(1).

### 2.4 Cells Coloring

A key element in the type of evolutionary experiments we model and study here, is to introduce an artificial color feature inherited through all divisions and mutations without altering major cell properties. In many evolutionary experiments and studies (see e.g. [18, 13, 10, 5, 28]) this binary coloring is implemented by pairs of inheritable biomarkers which do not alter cells biology. The initial white and red coloring generates two sub-populations (i.e. white and red) with independent evolutionary dynamics and naturally doubles the number of frequency variables to 2*g* in the model, since on each day *t* we then have *g* genotype frequencies among white cells and another vector of *g* genotype frequencies among red cells. The model presented above for the single-color evolution can be immediately extended to the two-colors evolution by doubling the number of frequencies in the frequency vector *h*. Therefore, in the two-colors stochastic model, for each day *t* and each *i* ∈ {1,2,…,*g*} we denote 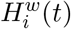 and 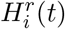 the respective frequencies of genotype-*i* in the white and red populations. Define then 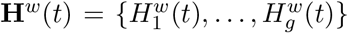 and 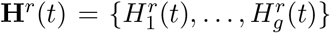. The state of the whole population is then described by the 2*g* dimensional vector of genotype frequencies **H**(*t*) = {**H**^*w*^(*t*), **H**^*r*^(*t*)}.

Because cells cannot change color by mutations, daily “growth+mutations” phases evolve independently for the white and red cells, and can hence be analyzed or simulated separately. But the daily selection step depends on the full “white + red” population after growth and mutations. Thus, formula (5) has to be modified appropriately to describe the random daily selection of *N* cells according to a multinomial distribution base on 2*g* frequencies *f_i_*, where *i* = {1,…, *g*} indexes the *g* white genotypes and *i* = {*g* + 1,…, 2*g*} indexes the *g* red genotypes.

## 3 Parameters Estimation of Stochastic Population dynamics

### 3.1 Formal Evolution Experiments

The type of genetic evolution experiments we consider here involves *n* independent populations evolving in parallel. This generates *k* independent evolutionary trajectories *pop^k^*(*t*), *k* = 1,…, *n*. Each population at time *t* = 0 starts from the same initial condition, namely having the same size *N*, containing only cells of genotype 1 (ancestor genotype), and colored 50/50 with red and white biomarkers.

Each population *pop^k^*(*t*) evolves in time and successive random emergence of beneficial mutations induce shifts in the relative proportion of white and red cells. Visibility of these shifts depends of course on the selective advantages of the emerging mutants. For each stochastic population trajectory *pop^k^*(*t*), there are the only two observables at each time *t* - the frequencies of red and white cells. This is because detailed genotype analysis is expensive and cannot be performed for each *t*. For each stochastic trajectory *pop^k^*(*t*), denote {*τ*_1_,*τ*_2_,…} the days of emergence of new mutants, and {*g*_1_,*g*_2_,…} as their respective genotypes. Of course, these two sequences of random variables are not directly observable.

Therefore, our goal here is to develop accurate estimators for the mutations emergence times {*τ*_1_, *τ*_2_,…} and the selective advantages {*s*_*g*_1__, *s*_*g*_2__,…} of the emerging mutant genotypes. We will focus on trajectories which have only a few mutations before overwhelming fixation of a strongly beneficial mutant. This is motivated by the Tim Cooper experiments on Escherichia Coli (see [28]) where most population trajectories exhibited a mutant fixation after one or two mutations. Trajectories with three or more mutations were fairly rare within 200 days runs, as could be also verified by intensive simulations. Moreover, for trajectories with 4 or more successive mutations, the analysis of later emerging mutants is a difficult task due to a high sensitivity of later mutations analysis to estimation errors on the parameters of earlier mutation events. Our simulations indicate that when a reasonably large number of observed trajectories are available, systematic analysis of the first two or three mutation events can provide fairly accurate estimates of the selective advantages of emerged mutants.

Some mutation events occurring during day *t* can remain unobservable due to the fact that new mutants can disappear during the random selection at the end of day *t*. However, some mutants strains emerging at the beginning of day *t* may grow fast enough during day *t* and survive the selection bottleneck at the end of day *t*. Thus, such mutants would grow even more on day *t* + 1 and affect the overall composition of the population (i.e. there will be a shift in the composition of white/red cells). Denote *τ*_1_ < *τ*_2_ < … the successive random times at which a new mutant strain emerges and persists for at least a few more days. For each observed population trajectory, we first implement an approximate estimation of times *T*_0_, *T*_1_, *T*_2_, *T*_3_, etc. which contain mutation events, i.e. *T*_0_ < *τ*_1_ < *T*_1_ < *τ*_2_ < *T*_2_ < *τ*_3_ < *T*_3_, etc. Next, we implement a non-linear least squares estimations of mutation times {*τ*_1_,*τ*_2_,…} and of the selective advantages during each mutation {*s*_*g*_1__, *s*_*g*_2__,…}. The main mathematical tool is the derivation of explicit analytical expressions for the most likely population histogram *h*(*t* + 1) given the (observed) histogram *h*(*t*).

## 4 Estimation of approximate intervals for the first and second mutation

First, we estimate approximate intervals for the first and second mutation events. This is carried out for each evolutionary trajectory and these intervals are then used in the optimization algorithm to compute the precise estimates for the first and second mutation times. Let *τ*_1_, *τ*_2_ be the emergence times of the first and second mutation events. Then our goal is to find *T*_0_ and *T*_1_ such that *T*_0_ < *τ*_1_ < *T*_1_ < *τ*_2_.

Let *w*(*t*) and *r*(*t*) = 1 − *w*(*t*) be the frequencies of white and red cells on day *t*, respectively. Since the white and red cells initially have the same genotype 1, this genetic composition persists for *t* ≤ *τ*_1_, so that one must observe *w*(*t*) ≈ *r*(*t*) ≈ 1/2 for *t ≤ τ*_1_ with fairly small random oscillations of *w*(*t*) and *r*(*t*) around 1/2. These small oscillations arise due to the random dilution step, since random selection of *N* cells after growth can alter slightly the proportion of white and red cells. For the Escherichia Coli experiments of *T*. Cooper [3] the empirical standard deviation *σ*_0_ of these oscillations can be crudely estimated on the first 30 days since the first beneficial detectable mutation events never occur before 30 days.

We define *T*_0_ as the last day *t* ≥ 1 such that |*w*(*u*) − 1/2| < 1.25 *σ*_0_ for all 1 ≤ *u ≤ t*. With high probability, the first beneficial mutation event occurs at some unknown time *τ*_1_ > *T*_0_. Then for *τ*_1_ < *t* ≤ *τ*_2_ one of the two frequencies *w*(*t*) or *r*(*t*) starts increasing at a faster rate than the other (i.e. |*w*(*t*) − 1/2| increases). Hence we define *T*_1_ to be the first day *t* such that |*w*(*u*) − 1/2| > 2*σ*_0_ for *u = t, t* + 1, *t* + 2 and all the (*w*(*u*) − 1/2) have the same sign. If (*w*(*u*) − 1/2) > 0 (or < 0) then we conclude that the first mutant strain emerges in the white population (or red population), respectively. With high probability we should have *T*_0_ < *τ*_1_ < *T*_1_, which we validated by numerical simulations outlined further on. As a test of our parameter estimation methodology, we used large sets of simulated stochastic genetic trajectories over time durations of 200 days. The simulated stochastic process is based on realistic mutation rates and selective advantages derived from the analysis of the *T*. Cooper experiments (see [3]). For our simulations, the frequency of the event *T*_0_ < *τ*_1_ < *T*_1_ < *τ*_2_ was of the order of 90% which is a fairly safe accuracy margin in our methodology, as can be seen below.

## 5 Estimation of parameters from the first mutant emergence

### 5.1 Descriptors of the first mutant emergence

As outlined in the previous section, for each observed population trajectory, our preliminary estimation algorithm analyzes the observed white cells frequencies *w*(*t*) to explicitly compute times *T*_0_ and *T*_1_ such that, with very high probability, one has *T*_0_ < *τ*_1_ < *T*_1_ < *τ*_2_. It is also easy to determine whether first mutant emergence occurs among white cells or among red cells. Next, we develop more precise estimation algorithm which utilizes observed white cells frequency data *w*(*t*) for all 0 ≤ *t* ≤ *T*_1_ to estimate the three unknowns *τ*_1_, *γ*_1_, *F*_*g*_1__ which defines a new genotype *g*_1_ > 1.

Denote *τ*_1_ the random day of emergence for the first new mutant strain which survives the selection phase on day *τ*_1_ and is, thus, present in the cell population at the beginning of day *τ*_1_ + 1. Denote *γ*_1_ > 0 the number of mutants emerging on day *τ*_1_ + 1. Let *g*_1_ be the genotype of these mutants, and denote their unknown growth factor as *F*_*g*_1__.

#### Notation

Define (*j, w*)-cells and (*j, r*)-cells as white and red cells of genotype *j*, respectively. Denote *w_j_*(*t*) and *r_j_*(*t*) the respective frequencies of (*j, w*)-cells and (*j, r*)-cells at the beginning of day *t*. Then, for 0 ≤ *t* ≤ *τ*_1_ all cells have the same ancestor genotype 1 so that *w_j_*(*t*) = *r_j_*(*t*) = 0 for *j* ≥ 2 and *w*_1_(*t*) = *w*(*t*), *r*_1_(*t*) = *r*(*t*). Recall that our computation of *T*_1_, also estimates whether the emerging mutants are white or red cells. These two cases are clearly similar, so without loss of generality we consider that the first emerging mutants are white cells. Analogous formulas can then be immediately derived for the case when emerging mutants are red cells. To simplify notation we drop the subscript as denote *τ* ≡ *τ*_1_ and *γ* ≡ *γ*_1_, since it is clear that we’re discussing the first mutation event. Let *η* be the number of white cell mutants present in the terminal population at the end of day *τ* right before the day *τ* selection. Then η has a Poisson distribution with mean λ = *N_μ_ F*_1_*w*(*τ*).

### 5.2 Growth phase

Since mutants of genotype *g*_1_ emerge in the white sub-population on day *τ* and survive the dilution (selection) step, at the beginning of day (1 + *τ*) we have

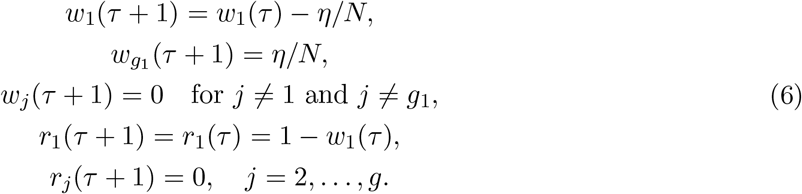

For *τ* + 1 ≤ *t < τ*_2_, new mutants may emerge during the day *t* mutation phase but they necessarily disappear during the day *t* selection phase because *t < τ*_2_. Since second mutation occurs on day *t* = *τ*_2_ and “survives” the dilution step, second mutation event affects population only on day *t* = *τ*_2_ + 1. Explicit analysis of cell frequencies on day *t* = *τ* + 1 is presented in (6). Then we consider days *t* > *τ* + 1 next. At beginning of day *t* with *τ* + 1 < *t < τ*_2_ the population contains only three cell types, namely (1, *w*)-cells, (*g*_1_, *w*)-cells, and (1, *r*)-cells, so that

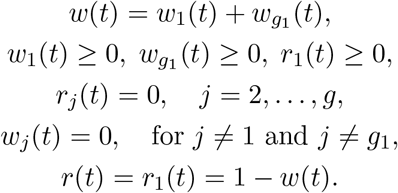

For *τ* + 1 < *t < τ*_2_, at the end of the day *t* growth phase, there are only three non-zero subpopulations with sizes

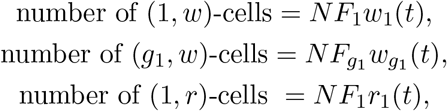

and all other sub-population sizes are zero. The total population size at the end of day *t* growth phase is *NK*, with *K* ≡ *K* (*t*) given by

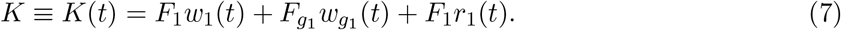

### 5.3 Mutations on day *t* for *τ* + 1 ≤ *t < τ*_2_

For days *τ* + 1 ≤ *t < τ*_2_ mutants of new genotypes can potentially emerge during the mutation step, but these new mutants are necessarily eliminated during the day *t* selection phase, since the second observable mutation occurs on day *τ*_2_. However, mutations of genotype 1 into genotype *g*_1_ can also occur and might influence the relative composition of white and red cells in the population. Thus, we analyze here the effect of mutations between genotype 1 and genotype *g*_1_.

Denote the random number of mutants generated at the end of day *t* by

*ν*_1,*j,w*_ = number of (1, *w*)-cells mutating into (*j, w*)-cells,
*ν*_*g*_1_,*j,w*_ = number of (*g*_1_, *w*)-cells mutating into (*j, w*)-cells,
*ν*_1,*j,r*_ = number of (1, *r*)-cells mutating into (j, r)-cells.

Recall that *ν*_1,1,*w*_ = *ν*_*g*_1_,*g*_1_,*w*_ = *ν*_1,1,*r*_ = 0. Moreover, *ν*_1,*j,w*_, *ν*_*g*_1_,*j,w*_, and *ν*_1,*j,r*_ are Poisson random variables with conditional means and variances as indicated in (2) so that

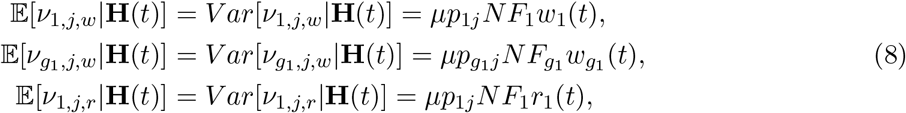

where **H**(*t*) denotes the total histogram of white and red cells. For *i, j* ∈ {1…*g*} denote *f_i,j,w_ = ν_i,j,w_/N* and *f_i,j,r_ = ν_i,j,r_/N*. Since *F_g_* is the largest growth factor, the standard deviations of *f_i,j,w_* and verify

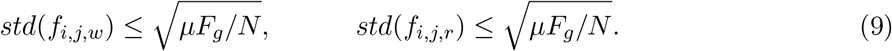

### 5.4 Multinomial Selection on day *t* for *τ*_1_ + 1 ≤ *t < τ*_2_

At the beginning of day *t* with *τ* + 1 ≤ *t < τ*_2_, there are only three non-zero sub-populations with frequencies

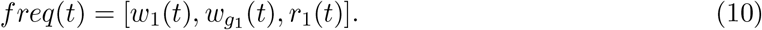

To preserve this property, the selection phase at the end of day *t* must realize the following event Ω(*t*) = {the random sample of size *N* selected at the end of day *t* contains only three cell types (1, *w*), (*g*_1_, *w*), (1, *r*)}. Therefore, if we denote the population after growth on day *t* as POP, then it is partitioned into three subpopulations POP(1, *w*), POP(*g*_1_, *w*), POP(1, *r*). We denote the population after the mutation step as MPOP. Denote *Mut*(*i, j, w*) the sets of (*i, w*) cells mutating into (*j, w*) cells, with a similar definition for red mutants *Mut*(*i, j, r*). Define

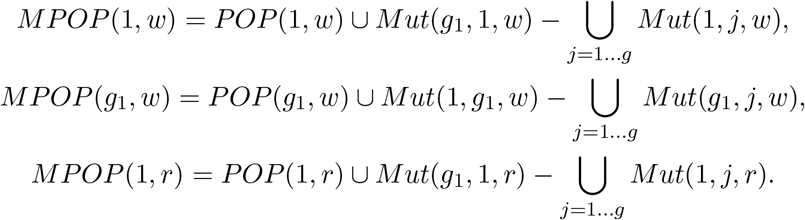

Define the Restricted Population *RestPop = MPOP*(1, *w*) ∪ *MPOP*(*g*_1_, *w*) ∪ *MPOP*(1, *r*). For *τ* + 1 ≤ *t < τ*_2_, the event Ω(*t*) is necessarily realized, and hence the random sample 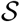 of size *N* selected at end of day *t* within MPOP *must be included* in RestPop. Therefore, given *freq*(*t*), the conditional distribution of genotypes within 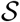 must be a *Multinomial*(*RestPop, N*), where one extracts a random sample of size *N* from the population RestPop. Therefore, changes in *POP* due to mutations can be quantified by

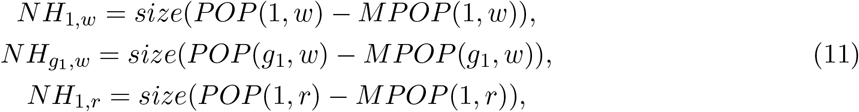

which implies

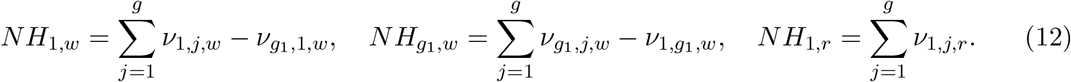

The size of *RestPop* is given by *size*(*RestPop*) = *N*(*K*(*t*) − *H*(*t*)), where

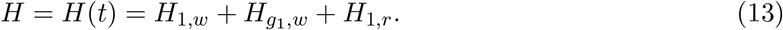

Due to (12) and (9), the sub-additivity of conditional standard deviation implies

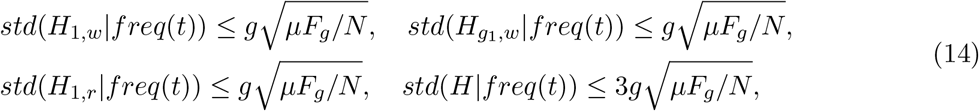

since *std*(*H*|*freq*(*t*)) = *std*(*H*_1,*w*_ + *H*_*g*_1_,*w*_ + *H*_1,*r*_{). The conditional means are given by

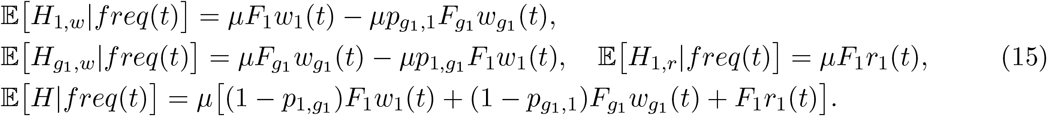

The first 3 conditional expectations are bounded by *μF_g_*, which in turn implies 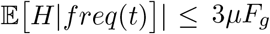. These bounds are of the order of the mutation rate with typical values *μ* ~ 10^−7^. Bounds for conditional standard deviations (14) are of the order of 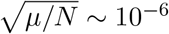 since *N* ≥ 10^5^.

Right before day *t* selection, frequencies for cell types (1, *w*), (*g*_1_, *w*), and (1, *r*) within the sub-population *RestPop* are given by

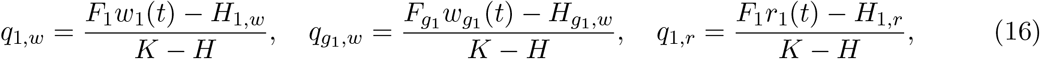

where *K* is the growth factor for the total population given by (7) and *H* is defined in (13). Since with high probability *H/K* = *O*(10^−6^), we introduce the approximation

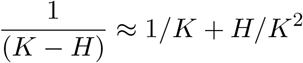

and, therefore, frequencies in (16) can be approximated as

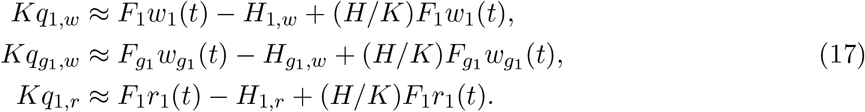

### 5.5 Recursive relation to predict *w*(*t* + 1) at time *t*, for *t* + 1 ≤ *t < τ*_2_

For *τ* + 1 ≤ *t < τ*_2_ the day *t* selection must realize the event Ω(*t*), which forces the day *t* multinomial selection to be restricted to the population *RestPop*. Given the vector *q* = [*q*_1,*w*_, *q*_*g*_1_,*w*_, *q*_1,*r*_] of nonzero frequencies in *RestPop*, the multinomial selection in *RestPop* entails

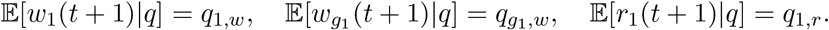

At time *t* with *τ* + 1 ≤ *t < τ*_2_, the best predictors *W*_1_(*t* + 1), *W*_*g*_1__(*t* + 1), *W*(*t* + 1) of *w*_1_(*t* + 1), *w*_*g*_1__(*t* + 1), *w*(*t* + 1) are defined by the conditional expectations

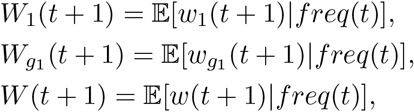

where *freq*(*t*) is defined in (10). In addition, we have the obvious relation *W*(*t* + 1) = *W*_1_(*t* + 1) + *W*_*g*_1__(*t* + 1). Since *K ≡ K*(*t*) is a deterministic function of *freq*(*t*), we then have

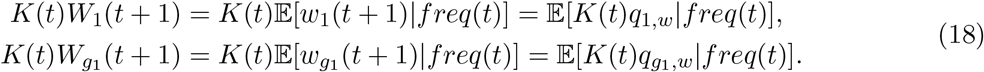

From (18) and (17) we then obtain

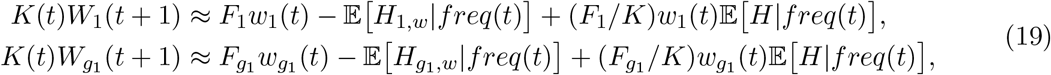

and due to (15), (19)

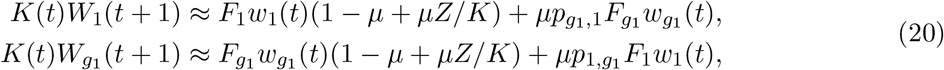

where

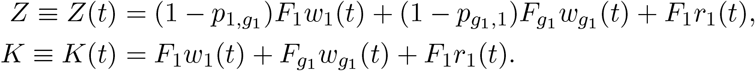

In equation (20) the only two unobserved freqencies are *w*_1_(*t*) and *w*_*g*_1__(*t*) = *w*(*t*) − *w*_1_(*t*), since *r*_1_(*t*) = *r*(*t*) is actually observed. The unobserved *w*_1_(*t*) can be approximated by its one-step conditional mean 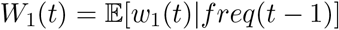, and *w*_*g*_1__(*t*) is then approximated by *w*(*t*) − *W*_1_(*t*). We thus obtain the following recurrence relation between *W*_1_(*t*) and *W*_1_(*t* + 1), as well as an evaluation of *W*_*g*_1__ (*t* + 1)

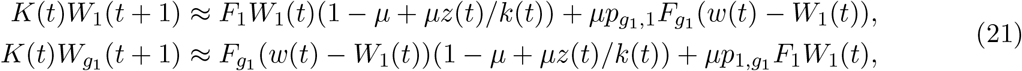

where *Z*(*t*) ≈ *z*(*t*) and *K*(*t*) ≈ *k*(*t*) with

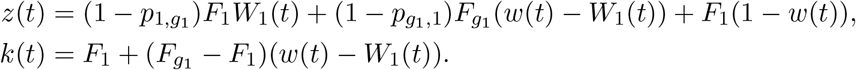

Therefore, The best predictor *W*(*t* + 1) of *w*(*t* + 1) verifies *W*(*t* + 1) = *W*_1_(*t* + 1) + *W*_*g*_1__(*t* + 1) and is hence given by

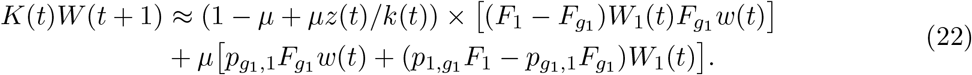

### 5.6 Non-linear least squares estimates of *τ*_1_, *γ*_1_, *s*_*g*_1__

As discussed in section 5.1, we first analyze each trajectory which contains daily frequencies of the white sub-population {*w*(*t*), *t* = 1, 2, …,} and estimate time-interval *RTIME* = [*T*_0_, *T*_1_] containing with very high probability the unknown emergence time *τ*_1_ of the first mutation.

In standard laboratory experiments for Escherichia Coli genetic evolution (see [3]), the growth factor *F*_1_ of the ancestor cells is well known, and the mutation rate *μ* can often be considered as already known from long term past experiments. One then seeks to estimate the selective advantage *s*_*g*_1__ of the first mutation event with unknown genotype *g*_1_. This selective advantage defines the growth factor *F*_*g*_1__ = (*F*_1_)^1+*s*_*g*_1__^. To develop an optimization procedure, one first needs to have an initial estimate for a plausible range for the unknown selective advantage. For instance, we take this range to be *RSEL* = [1%, 20%]. Moreover, we also need to have an estimate for a plausible range RMUT for the initial (unknown) number of new emerging mutants present at the beginning of day *τ*_1_ + 1. For Escherichia Coli experiments with *N* = 10^5^, one can for instance take *RMUT* = [1, 500].

We define an optimization procedure and perform a grid search over all possible combinations of (*τ, s, γ*) ∈ *RTIME × RSEL × RMUT*. We use the triplet (*τ, s, γ*) as a candidate for optimal values (*τ*_1_, *s*_*g*_1__, *γ*_1_) in formula (21) in the previous section. Given triplet (*τ, s, γ*) and white population *w*(*t*) at time *t*, the best predictor of the white population on the next day, *w*(*t* + 1), is 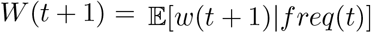 and the associated Squared Error is *SE* = (*w*(*t* + 1) − *W*(*t* + 1))^2^.

Recall, that *τ* is the estimator for the time of the first mutation event. Therefore, all cells have the same (ancestor) genotype until mutation and, thus, for *t* < *τ*

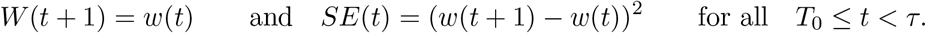

For *t = τ*, since *γ* mutants emerge on day *τ* + 1, we can approximate *W*(*τ* + 1) by

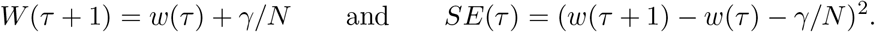

For *τ* + 1 ≤ *t ≤ T*_1_, we can recursively compute *W*(*t* + 1) from the first formula in (21), and this directly gives us the associated squared errors of prediction

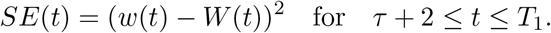

The Mean Squared Error is then computed as

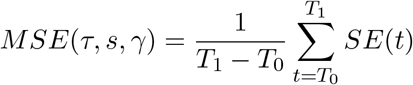

and the best estimate for the pair (*τ*_1_, *s*_*g*_1__) is computed by minimizing the MSE

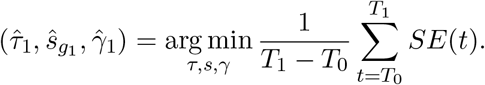

We solve the optimization procedure above by a straightforward multi-scale grid search over the set *RTIME × RSEL × RMUT*.

Theefore, *each observed trajectory* provides three estimates 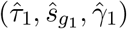 for the first mutants emergence time, for the selective advantage of these first mutants, and for the number of new mutants born on day *τ*_1_ and present at the beginning of day *τ*_1_ + 1, respectively. This optimization procedure also provides an estimate 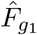 for the growth factor *F*_*g*_1__. In Section sec:pool we explain how we pool estimates for each individual trajectory into a global estimate over the whole ensemble.

## 6 Estimation of *T*_2_ such that 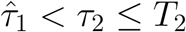

In previous sections we discussed how to use a single observed trajectory of white cell frequencies *w*(*t*) to estimate parameters 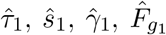 describing the first mutation event. After computing these estimates, we can use equations (21) iteratively to compute estimates for the evolution of the white sub-population until the second mutation event, i.e. we can use (21) to compute *W*_1_(*t*) and *W*_*g*1_ (*t*) of *w*_1_(*t*) and *w*_*g*1_ (*t*) for 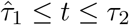.

Triple 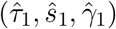 can be used to compute the behavior of mutants and estimate the frequency of the white sub-population until the second mutation. Estimates for the white sub-population are given by *W*(*t*) = *W*_1_(*t*) + *W*_*g*1_(*t*) for 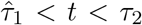 (here *τ*_2_ is unknown). On the other hand, observational data for the frequency of the white sub-population is also available, i.e. we observe *w*(*t*) for 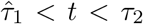. Therefore, we can compute the accuracy of prediction for *τ*_1_ < *t* < *τ*_2_ as *SE*(*t*) = (*w*(*t* + 1) – *W*(*t* + 1))^2^ and define the moving mean squared error of prediction

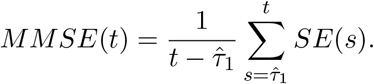

It is reasonable to assume that *MMSE*(*t*) remains reasonably small until the second mutation occurs, i.e. for *t* < *τ*_2_, but increases rapidly for *t* > *τ*_2_ (after the second mutation). This allows us to find an interval for the second mutation. In particular, we define *T*_2_ to be the first day 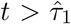 such that for *t* ≤ *u* ≤ *t* + 2 one has *SE*(*u*) > 2.5*MMSE*(*t*) and (*w*(*u*) – *W*(*u*)) has the same sign for *u* = *t*, *t* + 1, and *t* + 2. If the sign of *w*(*t*) – *W*(*t*) is positive (negative), then the second strain of new mutants emerges in the white (red) sub-population, respectively. We can also conclude that with high probability the second mutation occurs in the time interval 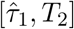.

## 7 Non-Linear least squares applied to the second mutant emergence

In section 5 we discussed to how to estimate parameters *τ*_1_, *γ*_1_, *s*_*g*1_ from the first mutation event. This also allows to compute the corresponding growth factor, *F*_*g*1_, of the new strain. Without loss of generality we can assume that the first mutation event occurs in the white sub-population. However, the second mutation event might occur in the sub-population of the same (white) or opposite (red) color and we have to analyze both cases. Denote *τ*_2_ the time of the second mutation event, *γ*_2_ the number of new mutants present at beginning of day *τ*_2_ + 1, and *g*_2_ the genotype of new mutants. The probabilistic analysis of the second mutant emergence is quite similar to the analysis outlined in section 5.

### 7.1 Case 1: the first and second mutation events occur in sub-population of the same color

Without loss of generality we assume that both the first and second mutant strains emerge among white cells. Thus (if *τ*_3_ is the time of the third mutation or the end of experiment), for *τ*_2_ < *t* < *τ*_3_, at beginning of day *t*, the only non-zero genotype frequencies are {*w*_1_(*t*), *w*_*g*1_ (*t*), *w*_*g*2_ (*t*), *r*_1_(*t*)}. Hence we have

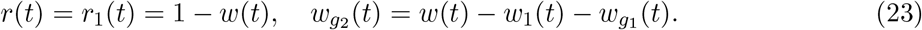

Therefore we define vector

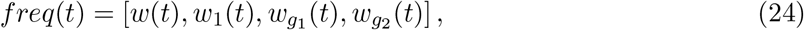

which contains all the information about frequencies of four sub-populations on day *t*.

Given *freq*(*t*), appropriate predictors for frequencies of white sub-populations *w*_1_(*t*+1), *w*_*g*1_ (*t* + 1), *w*_*g*2_ (*t* + 1), are given by

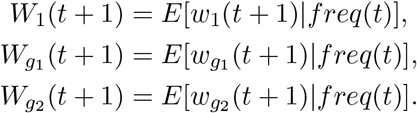

Similar to section 5.5, our goal is to obtain explicit formulas for conditional predictors for white frequency in the population. Thus, we can apply the same probabilistic arguments as in section 5.5 to derive the following formulas for frequencies of white sub-populations for all three genotypes. In particular, we obtain

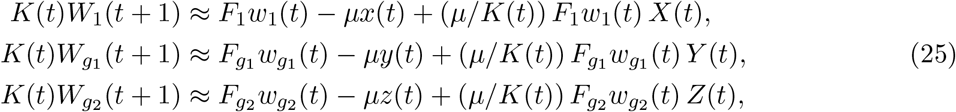

where

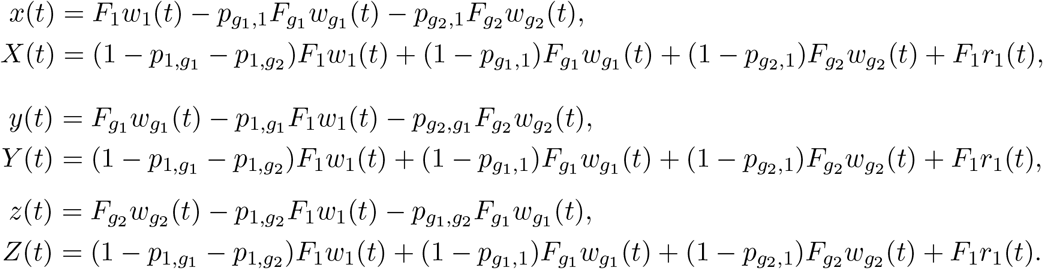

The term *K*(*t*) in equations (25) is given by

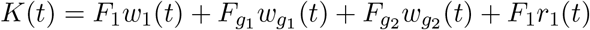

and due to the linear relations (23), *K*(*t*) can be expressed as

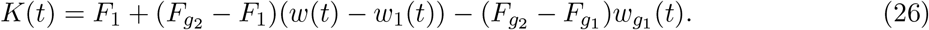

Formulas (25) and (26) define an explicit vector valued function Φ : *R*^3^ → *R*^3^ such that

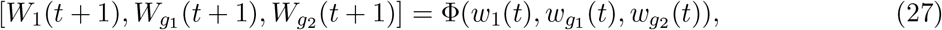

providing optimal predictions for frequencies of white sub-populations on day *t* + 1 given those frequencies on day *t*.

Since the second mutation occurs on day *t* = *τ*_2_ and becomes “visible” on day *t* = *τ*_2_ + 1, for 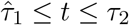, we have already computed (see section 5.5) estimates *W*_1_(*t*) and *W*_*g*1_(*t*) of frequencies of white sub-populations (1, *w*) and (*g*_1_, *w*). Combining two sub-populations provides an estimate *W*(*t*) = *W*_1_(*t*) + *W*_*g*1_ (*t*) for the frequency of white cells for 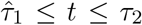. Since second mutation occurs at *t* = *τ*_2_, we assume that *w*_*g*2_(*τ*_2_) = 0, and substituting *W*_1_ (*τ*_2_) and *W*_*g*1_(*τ*_2_) into (27) to obtain estimates for *W*_1_(*τ*_2_ + 1), *W*_*g*1_(*τ*_2_ + 1). Next, we assume that *γ*_2_ mutants emerge during the mutation and, thus, the optimal estimate for the frequency of (*g*_2_, *w*) cells on day *τ*_2_ + 1 is *W*_*g*2_(*τ*_2_ + 1) = *γ*_2_/*N*. After that, for 2 + *τ*_2_ ≤ *t* ≤ *T*_2_ we expect our estimators *W*_1_(*t*), *W*_*g*1_(*t*), *W*_*g*2_(*t*) to be quite close to frequencies of (1, *w*), (*g*_1_, *w*), and (*g***2**, *w*) sub-populations since no mutations occur during that time-interval. Therefore, we can compute optimal estimates for frequencies of white sub-populations for the three genotypes iteratively by the recurrence formula

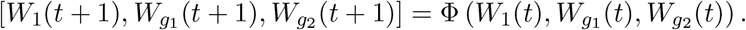

Combining estimates above, we obtain estimates for the frequency of white cells in the population *W*(*t*) = *W*_1_(*t*) + *W*_*g*1_(*t*) + *W*_*g*2_(*t*) for *τ*_2_ + 1 ≤ *t* ≤ *T*_2_. Combing the above expression with predictor *W*(*t*) for 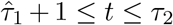 derived in section 5.5, we obtain explicit formulas for computing the optimal predictor of white-cell frequency *W*(*t*) for 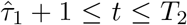. Therefore, the associated mean squared error of prediction can be computed as

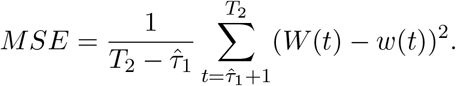

Of course the MSE above depends on the three unknown values *τ*_2_, *s*_*g*2_, *γ*_2_, which we want to estimate. Similar to discussion in section 5.6, we can define a non-linear least squares approach which consists in minimizing *MSE*(*τ*_2_,*s*_*g*2_,*γ*_2_) over all triplets (*τ*_2_,*s*_*g*2_,*γ*_2_) where *s*_*g*2_ ∈ *RSEL,γ*_2_ ∈ *RMUT* and 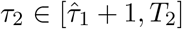. Similar to section 5.6 this minimization is done by multigrid search and is reasonably fast for the case of four genotypes. The minimizing triplet provides the least squares estimates 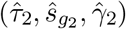.

### 7.2 Case 2: the first and second mutation events occur in sub-populations of distinct colors

In this section we derive formulas for optimal prediction of white- and red-cell frequencies when the first and second mutations occur in sub-populations of different colors. Since the approach used here is analogous to the discussion in sections 5.5 and 7.1, we only present the final result.

Denote conditional expectations of *w*_1_(*t* + 1), *W*_*g*1_ (*t* + 1), *r*_1_ (*t* + 1), *r*_*g*2_ (*t* + 1) given frequencies [*w*_1_(*t*),*W*_*g*1_(*t*), *r*_1_(*t*), *r*_*g*2_(*t*)] as *W*_1_(*t* + 1), *W*_*g*1_(*t* + 1), *R*_1_(*t* + 1), *R*_*g*2_(*t* + 1). Thus, we obtain the following four formulas valid for *τ*_1_ + 1 ≤ *t* ≤ *T*_2_

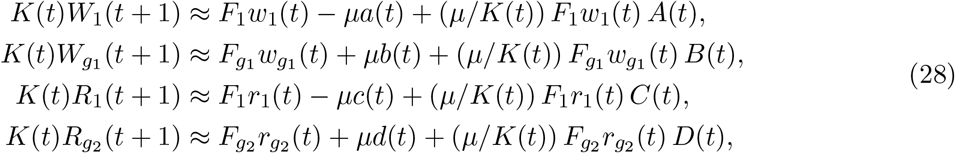

where

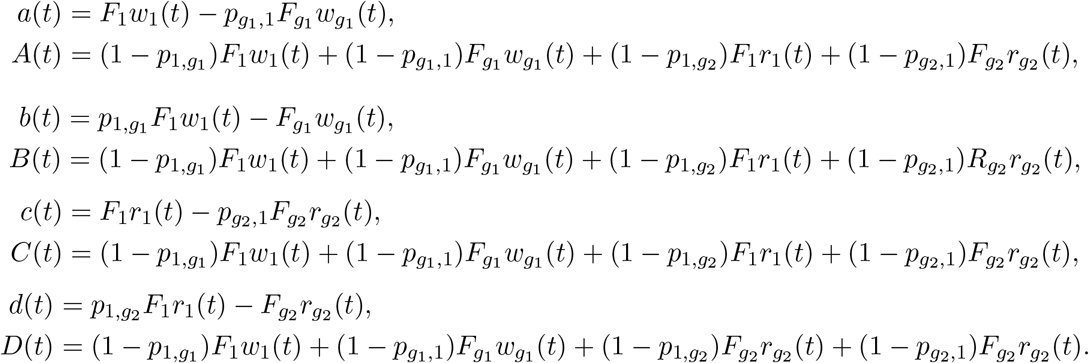

The term *K*(*t*) in equations (28) is given by *K*(*t*) = *F*_1_ *w*_1_(*t*) + *F*_*g*1_ *w*_*g*1_ (*t*) + *F*_1_*r*_1_(*t*) + *F*_*g*2_*r*_*g*2_ (*t*) and since we have *w*_*g*1_(*t*) = *w*(*t*) – *w*_1_ (*t*) and *r*_*g*2_(*t*) = *r*(*t*) – *r*_1_(*t*) = (1 – *w*(*t*)) — *r*_1_(*t*), *K*(*t*) becomes

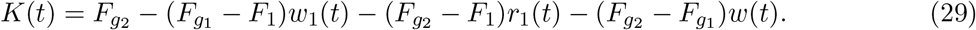

Similar to section 7.1 we can apply equations (28) to compute iteratively values of *W***1**(*t*), *W*_*g*1_ (*t*), *R*_1_(*t*), *R*_*g*2_ (*t*), for *τ***1** ≤ *t* ≤ *T*_2_. Since these terms are optimal predictors of white- and red-cells frequencies for different genotypes, the optimal predictor for the total frequency of the white-cell sub-population can be computed as *W*(*t*) = *W*_1_(*t*) + *W*_*g*1_ (*t*). Note, that the frequency of the white cells depends on the frequency of (*g*_2_, *r*) cells, *R*_*g*2_, through *A*(*t*) and *B*(*t*) defined above. Therefore, similar to section 7.1, we, can define the mean squared error and the corresponding optimization procedure to compute the estimates 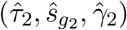.

## 8 Pooling estimates for the selective advantages of mutants

### 8.1 Multiple observed trajectories

In a typical evolutionary experiment, many colonies of the same bacteria (starting with the same ancestor genotype) evolve in parallel in a “rack” of distinct wells. All colonies undergo similar daily cycles of growth-mutations-selection. However, mutations do not occur in these colonies simultaneously. Therefore, such a multi-well long term experiment generates a finite number of different observed trajectories for white cell frequencies *w*(*t*). While all colonies start with the same ancestor genotype, white-cell frequencies for later times might differ for different colonies depending on a particular mutation which occurred in each colony. Observed long trajectories which exhibit zero emergence of new mutants strain generally constitute a small percentage of trajectories. As outlined above, we can automatically detect trajectories *ω*_1_,…,*ω_n_* having at least one emergence of a new mutant strain carrying a beneficial mutation. Next, for each observed trajectory *ω_j_*, we can apply our non-linear least squares estimation algorithm to generate an estimate 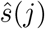 for the selective advantage of the first mutation event. The next step is to *combine these n estimates*. To this end, we treat individual estimates 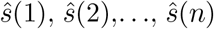 as independent observations and combine them into a histogram *H*_1_. Since in our model genotypes with higher selective advantage constitute a finite list of size *g* – 1, we expect histogram *H*_1_ to exhibit at most *g* – 1 local peaks centered around (unknown) *s*_2_,…, *s_g_*.

Of course we also apply our non-linear least squares analysis to the second emergence of a new mutant strain. This is done for *m* ≤ *n* trajectories exhibiting at least two new mutant strains, and provides another histogram *H*_2_ of *m* estimates of selective advantages. Similar to the histogram *H*_1_ discussed above, histogram *H*_2_ should exhibit at most *g* – 1 local peaks centered around *s*_2_,…, *s*_g_.

We have avoided mixing blindly the two histograms *H*_1_ and *H***2** because the frequency peaks in *H*_2_ are generally wider and hence less precisely centered than the peaks in *H*_1_. Thus, it is more efficient to first compute the center and dispersion of each peak in *H*_1_ to generate first estimators of *s*_2_, *s*_3_,…, *s_g_*. These estimators and their accuracy can then be boosted by a similar detection of new frequency peaks present in *H*_2_. This process can be extended to the analysis of the third emergence of mutant strains, to generate a histogram *H*_3_ of selective advantages estimates, and so on. However, the accuracy of frequency peaks in *H*_3_ is typically weaker than in *H*_2_. One of the reasons for this is that the number of trajectories exhibiting three successive mutation events is typically much smaller than *m*. Therefore, when we tested this approach on simulated trajectories, we restricted the histogram analysis to the first two mutations and histograms *H*_1_ and *H*_2_.

### 8.2 Gaussian Mixture Fitting to detect Histogram Peaks

The histogram *H*_1_ of *n* selective advantages described in the previous section typically has *g* – 1 peaks of frequencies. To detect them we fit a mixture of *g* – 1 Gaussians to this histogram. In particular, we consider a density function of the form

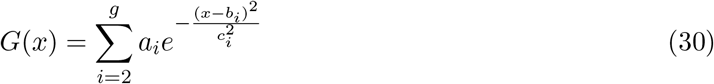

where *α_i_, b_i_, c_i_* are unknown positive parameters with the restriction 0 < *b*_2_ < *b*_3_ < … < *b_g_* since *b_i_* are the estimates for the selective advantages. Thus, there are 3(*g* – 1) parameters to optimize.

We fit *G*(*x*) to the histogram *H*_1_ by a standard non-linear least squares algorithm. This technique provides an acceptable fit when the number of parameters verifies 3(*g* – 1) ≪ *n*. Least squares fitting provides estimates 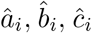 with theoretical accuracy of the order of 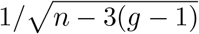. The *g* – 1 estimated peak centers 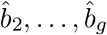 are then our estimates 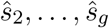 of the unknown selective advantages. Each estimated standard deviation 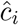 provides an approximate measure of the error for the corresponding estimate 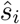.

In practice, the number of genotypes *g* is not really known a priori, and the analysis of histogram *H*_1_ enables a first estimation of *g* as the number of significant estimated peaks in *H*_1_. Moreover, analysis of the histogram *H*_1_ provides a set of estimates for the selective advantages of these genotypes. Subsequent analysis of the second histograms *H*_2_ provides a second set of estimates for the number of genotypes and their selective advantages. A statistical comparison between the first (based on *H*_1_) the second (based on *H*_2_) sets of estimates can then determine whether *H*_2_ has peaks significantly different from the *H*_1_ peaks, and improve (or validate) the estimation of *g*. After this comparison, the pairs of peaks in *H*_1_ and *H*_2_ which have significantly matching centers can then be combined to provide a more accurate estimation of selective advantages.

## 9 Results

In this paper we focus on validating the applicability and accuracy of the estimation approach using large synthetic datasets of simulated trajectories. In particular, we consider a model where every mutation occurrence in the population is equally likely to generate any one of the stronger mutants. In this case, the transition matrix 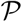 becomes

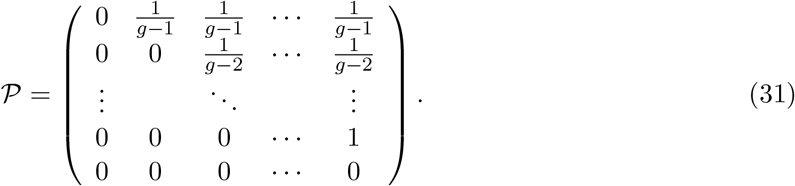

Therefore, entries of the matrix 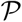 in our model are given by

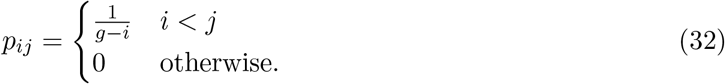

To validate our approach we generate a fixed number of trajectories using the above model with particular value of the mutation rate, *μ*, and selective advantages *s*_2_, *s*_3_, …, *s_g_*. We then apply our estimation algorithm to these simulated trajectories to recover estimates for the selected advantages 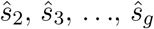.

### 9.1 Parameters of the simulated process

Parameters of the stochastic model for population growth are (i) the number of genotypes is *g* = 4; (ii) the number of simulated trajectories is *n* = 1000; (iii) *N* = 50000 is the daily initial population size; (iv) on the first day all cells have the same ancestor genotype with growth factor *F*_1_ = 200; (v) on the first day *N*/2 cells are white cells and *N*/2 cells are red cells; (vi) cell colors are preserved during growth, mutation, and selection; (vii) mutant genotypes have selective advantages *s*_2_ = 0.04, *s*_3_ = 0.07, *s*_4_ = 0.13; (viii) the mutation rate is *μ* = 10^−6^.

Selective advantages above correspond to growth factors (*F*_2_,*F*_3_,*F*_4_) = (247, 290, 398). These parameter values are consistent with numerical values reported in past and ongoing experiments for Escherichia Coli (see e.g. [9, 28]. We also fix the mutation transition matrix, as indicated in (32) which stipulates that only beneficial mutations can occur.

We simulated *n* = 1000 random trajectories with uniform length *T* = 200 days. At the beginning of each day *t* ∈ [1, 200], we record the total white cells frequency *w*(*t*) as well as the eight frequencies *w_j_* (*t*), *r_j_*(*t*), *j* = 1,…, 4 corresponding to sub-populations *j*-white or *j*-red cells. Of course, these detailed eight frequencies are not observable in actual experimental data, where only total frequencies of the white and red sub-populations are observed. Therefore, the eight unobservable frequencies are not used in computations of estimates for selective advantages. Nevertheless, frequencies of white and red sub-populations for each genotype provide additional data for validating our approach and, in particular, for validating estimation of intervals for the first and second mutations discussed in sections 4 and 6, respectively.

Examples of trajectories with one mutation and two mutations are presented in Figures 1 and 2, respectively. We consider that a trajectory reaches *fixation* at time *T_end_* if either *w*(*T_end_*) ≥ 95% or *w*(*T_end_*) ≤ 5% and our Figures thus involve only the days *t* ≤ *T_end_*. For brevity we call *mutation event* the emergence of a new mutant strain before fixation time.

**Figure 1:**
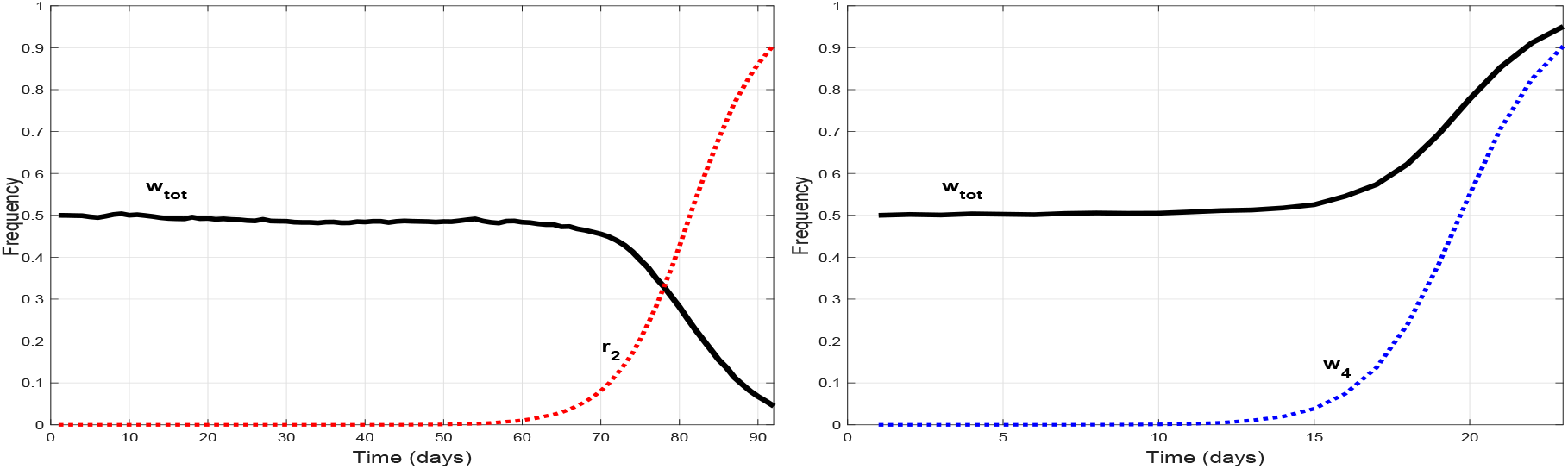
Two examples of trajectories with only one mutation event. We display the frequencies of white cells and of the sub-population of mutants. Black line (labeled *w_tot_*) = total white cells frequency *w*(*t*), Left part - emergence of genotype-2 mutants among red cells (red line *r*_2_), Right part - emergence of genotype-4 mutants among white cells (blue line *w*_4_).

**Figure 2:**
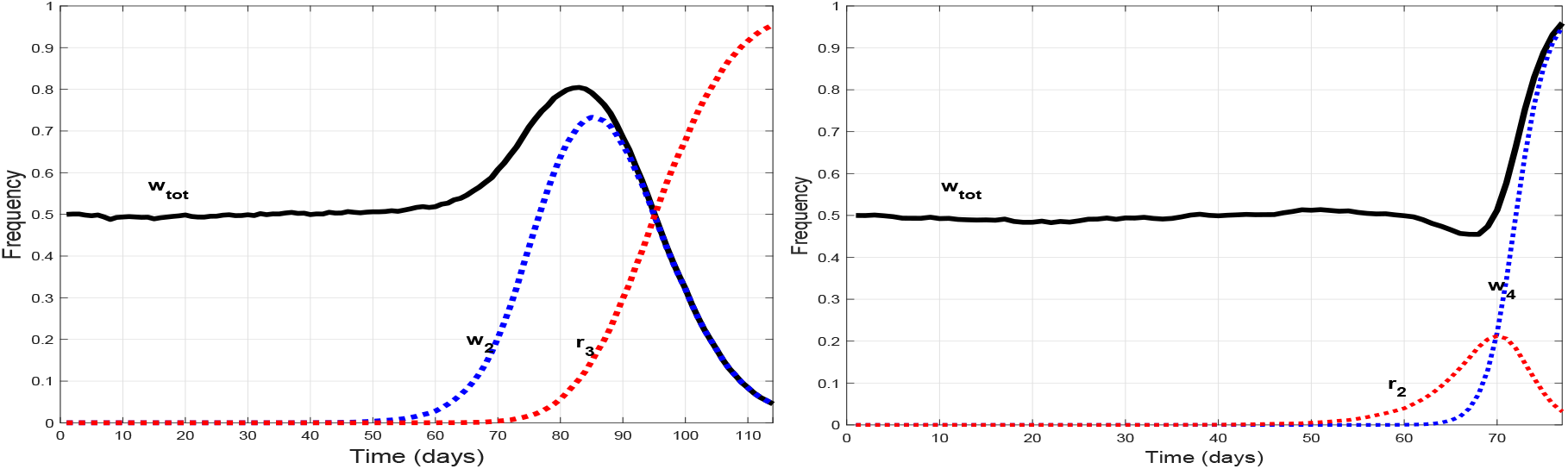
Two examples of trajectories with two mutations. We display the frequencies of white cells and of sub-populations of mutants. Black line (labeled *w_tot_*) = total white cells frequency *w*(*t*), Left part - emergence of genotype-2 mutants among white cells (blue line *w*_2_) and later emergence of genotype-3 mutants among red cells (red line *r*_3_), Right part - emergence of genotype-2 mutants among red cells (red line *r*_2_) and later emergence of genotype-4 mutants among white cells (blue line *w*_4_).

We considered that a mutation of type *j*-white had actually emerged at time *τ* < *T_end_*, if *τ* is the smallest time *t* such that *w_j_* (*t*) ≥ 1/1000. Same convention for mutations of type *j*-red cells. Among the *n* = 1000 simulated trajectories of duration *T* = 200 days 0.5% trajectories have no mutation event, 70% trajectories have only one mutation event, 23% trajectories have two mutation events, and 6.5% trajectories have three or more mutation events. The mean fixation time for trajectories with exactly *k* mutation events increases with k, starting from *T_end_* ≈ 70 for *k* = 1 and reaches the value *T_end_* ≈ 180 for *k* = 3. We also simulated a much larger number of trajectories *n* = 100, 000 to estimate the frequency of emergence *f*(*j*) for mutant genotypes *j* = 2,…, 4. This yielded *f*(2) = 19%, *f*(3) = 29%, *f*(4) = 52%. This implies that more mutation events is recorded for stronger genotypes and, therefore, the error of estimating selective advantages would decrease for mutant genotypes with larger fitnesses (bigger *j*).

### 9.2 Selective Advantages Estimation based on 1,000 trajectories

To validate our estimation approach we use only data for the total frequency of white cells, *w*(*t*), and we first estimate approximate time of the first mutation (see section 4) for each trajectory. Out of *n* = 1000 simulated trajectories, 995 trajectories exhibit at least one mutation event. Thus, we apply our least-squares algorithm to these 995 trajectories to estimate selective advantages of mutant genotypes (section 5.6) and then combine these estimates into the histogram *H*_1_ (section 8).

Histogram *H*_1_ and the corresponding fit *G*(*x*) in (30) are depicted in Figure 3. Histogram *H*_1_ has clear three local maxima with corresponding estimates for selective advantages 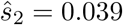, 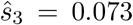, and 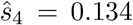. These estimation results are in a very good agreement with true parameter values *s*_2_ = 0.04, *s*_3_ = 0.07, and *s*_4_ = 0.13. Estimation errors for all three parameters are less than 5%. Standard deviations for the multi-Gaussian fit (*c*_2_,*c*_3_, *c*_4_) ≈ (0.012, 0.006, 0.005), provide estimates for the relative accuracy of estimation of selective advantages. Here we can clearly see that selective advantages of the two stronger phenotypes are estimated more accurately than the selective advantage of the two weaker mutant phenotype.

**Figure 3:**
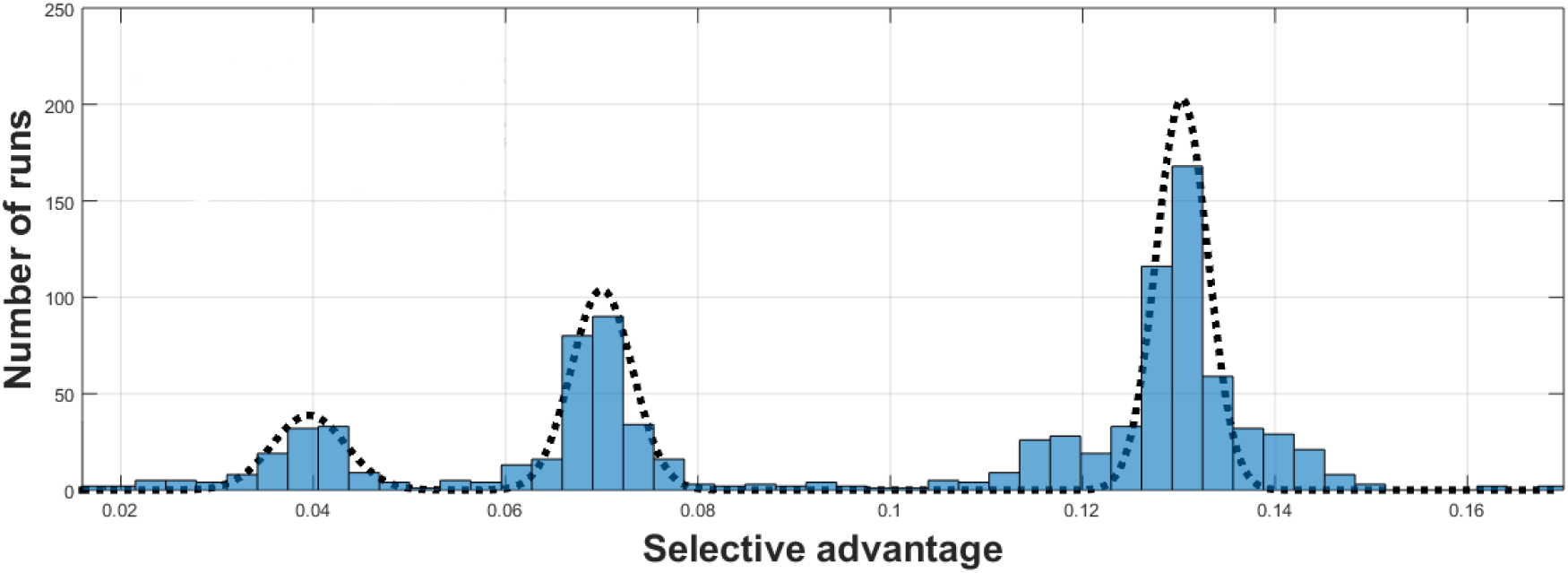
Histogram of the 3 selective advantages based on the first mutation event for 995 simulated trajectories and corresponding fit *G*(*x*) in (30). Corresponding estimates for the selective advantages are *ŝ*_2_ = 0.039, *ŝ*_3_ = 0.073, and *ŝ*_4_ = 0.134.

In addition to the analysis of the first mutation event, we also analyzed the emergence of second mutation in 995 trajectories as discussed in sections 6 and 7. Approximately 300 trajectories exhibit a second mutation event and for each one of those trajectories, we applied our non-linear least squares algorithms to analyze the second emergence of the second mutant strain. This generated a histogram *H*_2_. However, histogram *H*_2_ in this case has only one clearly pronounced peak. Moreover, multi-Gaussian fit for the histogram *H*_2_ resulted in much larger (at least one order of magnitude) standard deviations for the standard deviations *c_j_, j* = 2,…, 4. Therefore, we concluded that 300 trajectories are not sufficient to accurately estimate selective advantages based on the second mutation event and combine histograms *H*_1_ and *H*_2_.

### 9.3 Selective Advantages Estimation based on 10,000 trajectories

To evaluate more precisely the impact of the second mutation event analysis on selective advantages estimates, we also performed estimation of selective advantages from *n* = 10,000 trajectories with duration 200 days each. This resulted in approximately 3700 trajectories exhibiting at least 2 mutation events.

In figure 4, histograms *H*_1_ (top) and *H*_2_ (middle) depict results of estimation based on the first and second mutation events, respectively. Histogram *H*_1>2_ is obtained by combining *H*_1_ and *H*_2_. There are much fewer trajectories exhibiting at least two mutation events compared with trajectories exhibiting at least one mutation event. Therefore, estimates based on the second mutation event have much larger standard errors than the *H*_1_ estimates. Although the number of trajectories exhibiting a second mutation event is relatively high, the combined histogram *H*_1,2_ provides considerably less accurate estimates for selective advantages, especially for the genotype *j* = 2. We would like to point out that these results are particular to the mutation matrix *P* in (31). In particular, for the mutation matrix *P* considered here all mutant genotypes (*j* > 1) can emerge directly from the ancestor genotype *j* = 1 with equal probabilities 1/(*g* − 1). Therefore, we can conjecture that in such cases analysis of the first mutation event contains sufficient information to adequately estimate selective advantages if there are many trajectories exhibiting the first mutation event. However, there are several situations where analysis of the second mutation event can provide useful information. We comment about this in the conclusions.

**Figure 4:**
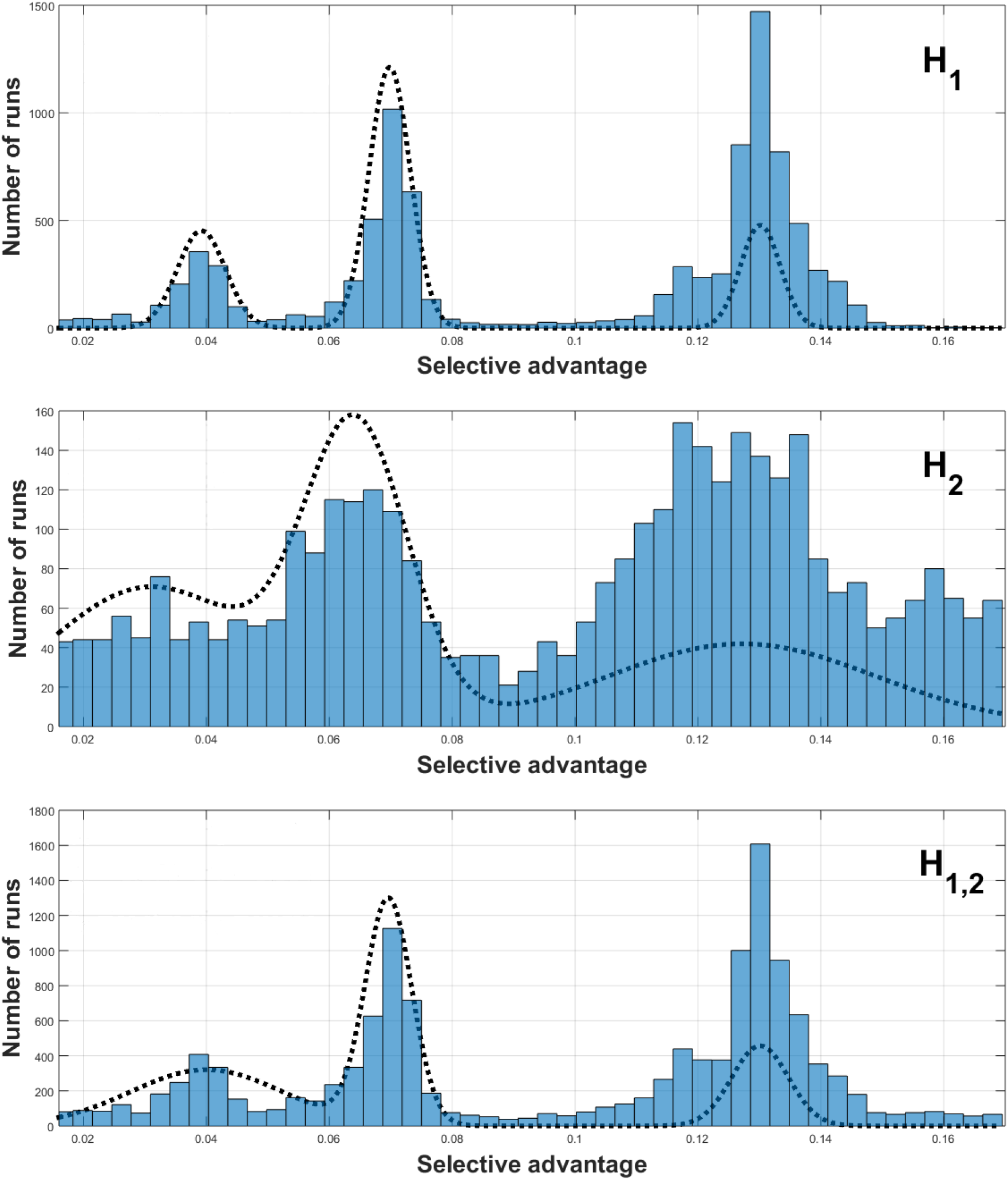
Gaussian mixture estimates for selective advantages. Results are based on algorithmic analysis of *n* = 10,000 simulated trajectories. Top : Estimates of selective advantages by analysis of first new mutant strain; Middle : Estimation of selective advantages by analysis of second new mutant strain; Bottom : Estimation of selective advantages by joint analysis of first and second new mutant strain.

We also analyzed the behavior of standard error of estimations for selective advantages as the number *n* of simulated trajectories increases. In particular, standard errors *c_j_*(*n*) with *j* > 1 were computed for *n* = 75, 150, 250, 1000, 10000, 50000, 100000. A linear regression demonstrated that for *n* ≥ 250 standard errors decay as *c_j_*(*n*) ~ *n*^−1/2^. It is possible to provide an analytical argument which supports this conclusion for large *n*.

## 10 Conclusions

In this paper we present an efficient non-linear least squares approach to estimate selective advantages of mutant genotypes, based on the type of observed data provided by long-term experiments focused on bacterial genetic evolution. In many such experiments, bacterial cells are colored with two biomarkers (e.g. white and red), and on each “day” *t* the frequency of white and red cells is recorded. We analyze a Markov chain model which corresponds to this experimental setup for evolutionary experiments. The dynamics of this model is based on daily cycles which include three phases - deterministic growth, random mutations, and random selection (dilution). The stochastic model involves a finite set of *g* distinct genotypes with one ancestor genotype and *g* − 1 mutant genotypes with unknown selective advantages.

Typically, evolutionary experiments involve n independent populations evolving in parallel, all starting with a population of *N* cells having the same ancestor genotype. Therefore, such evolutionary experiments usually generate an ensemble of time-series (trajectories) of observed white cells frequencies. For each trajectory, we perform least-squares estimation of selective advantages by minimizing the mean squared error between the observed white-cell frequency, *w*(*t*), and prediction of the Markov chain model, *W*(*t*). We then gather these n estimates of selective advantages over all observed trajectories to generate a histogram of estimated selective advantages. Typically, we expect this histogram to have several peaks which correspond to selective advantages of mutant genotypes. We then approximate the histogram by a multi-Gaussian mixture to estimate values of selective advantages. In addition, variances in the muti-Gaussian approximation provide rough estimate on the uncertainty in the estimation of the corresponding selective advantages.

We demonstrated applicability of our approach by applying it to synthetic datasets generated by the Markov chain model discussed above. We verified that our estimation approach is able to recover the selective advantages used in the model. As an outcome of our numerical investigation we can can conclude that our approach produces accurate results for ensembles sizes of 100 or larger. Our approach can be applied to smaller ensembles, but estimation errors can become quite large. In this paper we present numerical results with *g* = 4 genotypes. We also validated our approach for *g* = 6 genotypes. In addition, we also validated that errors of estimation decrease at speed proportional to the square root of the ensemble size, and that the estimates for selective advantages are reasonably accurate as soon as the ensemble size is bigger than 200.

Mathematical analysis of the evolutionary model presented in this paper enables estimation of selective advantages using the first mutation event or a combination of the first and second mutation events. Our numerical results demonstrate that a straightforward pooling of estimators obtained from the first and second mutation events may not be always optimal. Instead, analysis of the second mutation event can be used to extend results of the estimation from the first mutation event data. However, analysis of the second mutation event can be quite useful for improving (or even discovering) new mutant types and/or improving a subset of the selective advantages estimates. First, second mutation events can be used to identify genotypes which were rarely seen in first mutations events, but more frequently present among second mutation events. Such situations can occur for small ensemble sixes *n* ≈ 100,…, 200. Second, analysis of the second mutation event can shed light on the shape of the mutation matrix *P* and whether all mutant genotypes can emerge directly from the ancestor genotype. In other words, analysis of the second mutation event can potentially elucidate importance of double mutations. Finally we have also successfully tested that our approach can easily be extended to the estimation of the mutation rate *μ*. Additional simulations and parameter estimation studies can be found in [24].

Realistic evolutionary experiments include a rather small number of populations growing in parallel. However, these experiments are carried out for many years, and bacterial populations are often re-seeded with the original ancestor genotypes after a dominant mutation occurs and one color overtakes the whole population. Therefore, a realistic number of trajectories available for estimating selective advantages are of the order of 100. Nevertheless, estimation procedure described here is still applicable even with such small number of available trajectories.

For small number of trajectories our approach should then be applied sequentially as follows. First, compute and analyze histogram *H*_1_ derived from first mutation events and compute first estimates of selective advantages as well as their standard estimation errors. Second, repeat optimization procedure for computing estimates for selective advantages based on first mutation events (section 5.6) with improved search intervals for *s ∈ RSEL*. In particular, *RSEL* can be taken as union of confidence intervals for selective advantages computed on the first pass using multi-Gaussian fit of *H*_1_. We expect that the second pass of the optimization procedure based on the first mutation event data should yield much more accurate estimates for time of the first mutation, *τ*_1_, selective advantage *s*_*g*_1__, and the number of emerged mutants, *γ*_1_. In addition, second pass should also yield an improved histogram of selected advantages, *H*_1_. Third, perform optimization procedure for the second mutation event data with *s ∈ RSEL*, where *RSEL* has been computed from the improved *H*_1_. In contrast with the present paper where analysis of the second mutation event is based on the selective advantage of the first mutant genotype computed from a current (single) trajectory, using estimates from *H*_1_ is more robust and should provide a more accurate version of *H*_2_, with relatively small standard errors. The least squares fitting of the multi-Gaussian mixture for *H*_2_ should then be performed conditional on the information gathered by the *H*_1_ estimates and their standard errors. This approach will provide better estimates of selective advantages derived from *H*_2_. The fusion of estimates obtained from *H*_1_ and *H*_2_ should be performed on the basis of their respective standard errors. We plan to implement and test this upgraded approach in a subsequent paper, applying our estimation procedure to realistic observational data.

## Acknowledgment

This research has been partially supported by NSF grants DMS-1412927 and DMS-1903270.

